# Comparative analysis of mammal genomes unveils key genomic variability for human lifespan

**DOI:** 10.1101/2021.02.09.430384

**Authors:** X. Farré, R. Molina, F. Barteri, P.R.H.J. Timmers, P.K. Joshi, B. Oliva, S. Acosta, B. Esteve-Altava, A. Navarro, G. Muntané

## Abstract

Mammals vary 100-fold in their maximum lifespan. This enormous variation is the result of the adaptations of each species to their own biological trade-offs and ecological conditions. Comparative genomics studies have demonstrated that the genomic factors underlying the lifespans of species and the longevity of individuals are shared across the tree of life. Here, we set out to compare protein-coding regions across the mammalian phylogeny, aiming to detect individual amino acid changes shared by the most long-lived mammal species and genes whose rates of protein evolution correlate with longevity. We discovered a total of 2,737 amino acid changes in 2,004 genes that distinguish long- and short-lived mammals, significantly more than expected by chance (p=0.003). The detected genes belong to pathways involved in regulating lifespan, such as inflammatory response and hemostasis. Among them, a total 1,157 amino acids, located in 996 different genes, showed a significant association with maximum lifespan in a phylogenetically controlled test. Interestingly, most of the detected amino acids positions do not vary in extant human populations (>81.2%) or have allele frequencies below 1% (99.78%), Consequently, almost none could have been detected by Genome-Wide Association Studies (GWAS). Additionally, we identified four more genes whose rate of protein evolution correlated with longevity in mammals. Crucially, SNPs located in the detected genes explain a larger fraction of human lifespan heritability than expected by chance, successfully demonstrating for the first time that comparative genomics can be used to enhance the interpretation of human GWAS. Finally, we show that the human longevity-associated proteins coded by the detected genes are significantly more stable than the orthologous proteins from short-lived mammals, strongly suggesting that general protein stability is linked to increased lifespan.

## Introduction

Why some individuals within a species live longer than others is intimately related to the broader question of why some species live longer than others (Tian, Seluanov, and Gorbunova 2017). Maximum lifespan (MLS) is a species-specific trait: flies, dogs, and humans all have different but consistent lifespans that are adapted to their ecology and biology. From an evolutionary standpoint, the main ultimate cause of these differences are lineage-specific ecological adaptations that modify the rates of extrinsic mortality. For instance, lifespan is usually lengthened by arborealism (Shattuck and Williams 2010), flight (Pomeroy 1990), subterranean life (Buffenstein 2005), and body mass (Austad 2005; de Magalhães, Costa, and Church 2007) since all these adaptations reduce extrinsic mortality by predation. Most of these differences and similarities in MLS are explained by common physiological, biochemical, and genetic basis across species (Ma and Gladyshev 2017).

Mammals show a 100-fold variation in MLS, ranging from short-lived species like forest shrews (~2 years) to long-lived species like the bowhead whale (~200 years, Tacutu et al. 2018), representing an ideal lineage to study the genomics of lifespan and to unveil genes and pathways that may be relevant for humans. Numerous studies have been devoted to study mammal lifespan focusing on individual species, such as the bowhead whale (Keane et al. 2015) and the naked mole rat (Kim et al. 2011; Ruby, Smith, and Buffenstein 2018), or relatively small subgroups like bats (Huang et al. 2019; Wang et al. 2020; Seim et al. 2013). While single-species studies have yielded some credible candidate genes associated with increased lifespan, it is difficult to obtain generalizations on universal mechanisms of lifespan regulation from them. Therefore, knowledge about lifespan evolution in mammals is still limited. Our mammalian ancestors, although diverse (Pickrell 2019), were small (O’Leary et al. 2013), and in all likelihood short-lived creatures. Whereas the long-term directional bias towards increasing size in mammals (Baker et al. 2015; Lyson et al. 2019) could drive a parallel trend towards increased lifespan, there is no strong evidence in favor of that hypothesis.

Other comparative genomics studies have focused on identifying rapid evolutionary changes in genomes or transcriptomes that correlate with changes in longevity (Kim et al. 2011; Muntané et al. 2018; Kowalczyk et al. 2020); or have assessed the relationship between lifespan and other adaptations with life-history traits in different taxa (Montgomery and Mundy 2012; Boddy et al. 2017; Wang et al. 2020; Zhang et al. 2014; Chikina, Robinson, and Clark 2016; Foote et al. 2015). These studies have identified longevity pathways that are conserved across species, such as the insulin/IGF-1 pathway, telomere maintenance, DNA repair, coagulation and wound healing, proteostasis, and TOR signaling. The existence of such common pathways and mechanisms is consistent with the fact that long-lived animals show convergent phenotypes, including increased stress resistance, altered metabolism, and delayed reproduction and development (Hekimi 2003).

A key mechanism that may contribute to differences in lifespan is the maintenance of the proteostasis network. Protein stability or proteostasis refers to the capacity to protect protein structures and functions against environmental stressors, including aging. In fact, dysfunction of the protein quality control mechanisms is a hallmark of aging (López-Otín et al. 2013; Santra, Dill, and de Graff 2019) and there is substantial evidence linking proteostasis and longevity (reviewed in Tian, Seluanov, and Gorbunova 2017). For instance, improved protein stability is determinant for longevity in exceptionally long-lived mollusks (Treaster et al. 2014) and in the naked mole-rat, the longest-living rodent (Pérez et al. 2009). In addition, interventions that enhance proteome stability can improve health or increase lifespan in model organisms (Fontana and Partridge 2015), such as pharmacological chaperones that have been investigated as potential therapeutic targets to reduce the adverse effects of. misfolding of aging-related proteins (Powers et al. 2009; Bullock et al. 1997).

Despite of all the evidence outlined above, a mammalian-wide study of the genomic underpinnings of lifespan has never been carried out with the combined goals of identifying individual mutations linked to longevity; analyzing the functional properties of their genes and the pathways in which they take part; and studying how the stability of the proteins coded by these genes may differentiate long- and short- lived species. In fact, the largest-scale studies conducted on mammalian lifespan have focused only on humans, with somewhat limited results (e.g., Timmers et al. 2019). The low heritability of lifespan in humans may explain these limitations. Studies in twins have reported values of heritability between 0.2 and 0.3 (Herskind et al. 1996; Sebastiani and Perls 2012). More recently, the analysis of family trees produced an estimation of around 0.1, suggesting that previous estimates were inflated due to assortative mating (Kaplanis et al. 2018; Ruby et al. 2018). Several GWAS on human lifespan have been carried out using different indirect measures and strategies, such as parental lifespan (Timmers et al. 2019), extremely long lived individuals (Deelen et al. 2019), or health span (Zenin et al. 2019). While these studies have identified a set of genetic variants that are associated with an individual’s lifespan, only a small fraction of the heritability –around 5%– has been disclosed by GWAS. In summary, not only we are missing important contributions to extant human variation on lifespan, probably due to genetic variants with small effects (de Magalhães and Wang 2019; Muntané et al. 2018); but also, and given the relatively small genetic variation in lifespan in our species, we still lack a map of the full landscape of genomic factors underlying lifespan.

Here, we performed the largest phylogeny-based genome-phenome analysis to date, focusing on the detection of individual mutations and genes that underlie the enormous variation of lifespan in mammals. We report the discovery of more than 2,000 longevity-related genes and show that, overall, they present a trend towards increased protein stability in long-lived organisms. In addition, we successfully show that our findings enhance the interpretation of the results of lifespan GWAS that have been carried-out in humans. Altogether, our results pave the way for the use of comparative genomics studies to shed light on human traits, particularly those of potential medical interest.

## Materials and Methods

### Genomic and phenotypic data

Amino acid (AA) and nucleotide alignments for 39,178 orthologous coding sequences were retrieved from the Multiz alignment of 100 vertebrate genomes (Human 100-way) together with the mammalian phylogenetic tree, which was also downloaded from UCSC (https://genome.ucsc.edu/, last accessed August 2019). Amongst the 100 vertebrate species, we kept the 62 species belonging to the class Mammalia (Supplementary Figure 1). For each gene, only the longest transcript was kept and protein alignments with an overall number of gaps > 50% or in human alternate contigs were excluded (n=905). After this filtering, a total of 18,266 protein transcripts were included in the analyses.

Variation in MLS across species correlates with many life-history traits, including body mass, growth rate, age at sexual maturity, and body temperature, which can bias comparative studies of lifespan (Speakman 2005). The most relevant and studied confounding factor is body mass, so longevity is usually corrected by it using the longevity quotient (LQ), which indicates whether a species has an average lifespan or is unusually long- or short-lived relative to its body size. LQs is computed as the ratio of a species MLS to the expected MLS given its body mass (Austad and Fischer 1991). MLS and adult body mass were obtained from the AnAge database (Tacutu et al. 2018, build 14) and missing information was complemented, when available, using data from the Animal Diversity Web (Myers et al. 2019). The LQ of each species was calculated using the allometric equation for mammals (de Magalhães, Costa, and Church 2007). After filtering out species for those we were unable to obtain LQ data, we kept a final number of 57 mammalian species for subsequent analyses (Supplementary Figure 1 and Supplementary Table 1).

### Convergent Amino Acid Substitutions: Discovery and Validation

Convergent amino acid substitutions (CAAS) are AA changes that have occurred independently at least twice across the phylogeny. For the purposes of this work, we focused on CAAS that coincide with extreme lifespan values in the set of mammalian species under study. We designed a two-phase procedure to identify such instances of CAAS. First, in the *Discovery phase*, we selected the species in the top and low deciles of the LQ distribution, which we named long-lived and short-lived, respectively, for a total of 12 extremely lived species (6 top and 6 low). Subsequently, an in-house script was used to detect specific protein positions in which the reference genomes of the long-lived species had the same AA and the short-lived group presented either another AA (Scenario 1) or a set of segregating AAs that were different from the reference AAs in the long-lived-group (Scenario 2). Positions where the short-lived group showed a fixed AA, and where segregating, non-intersecting variation was observed in the long-lived group were also considered (Scenario 3). For the purposes of this work, we only focused on Scenarios 1 and 2, representing the AA substitutions converging in the mammal long-lived species (discussed in Supplementary Note). We required full information from all species in the extremes, so AA positions for which one or more of the species had a gap were excluded from the analysis. Such a filter resulted in a final set of 13,035 genes evaluated using the CAAS procedure (Figure 1).

**Figure 1.**
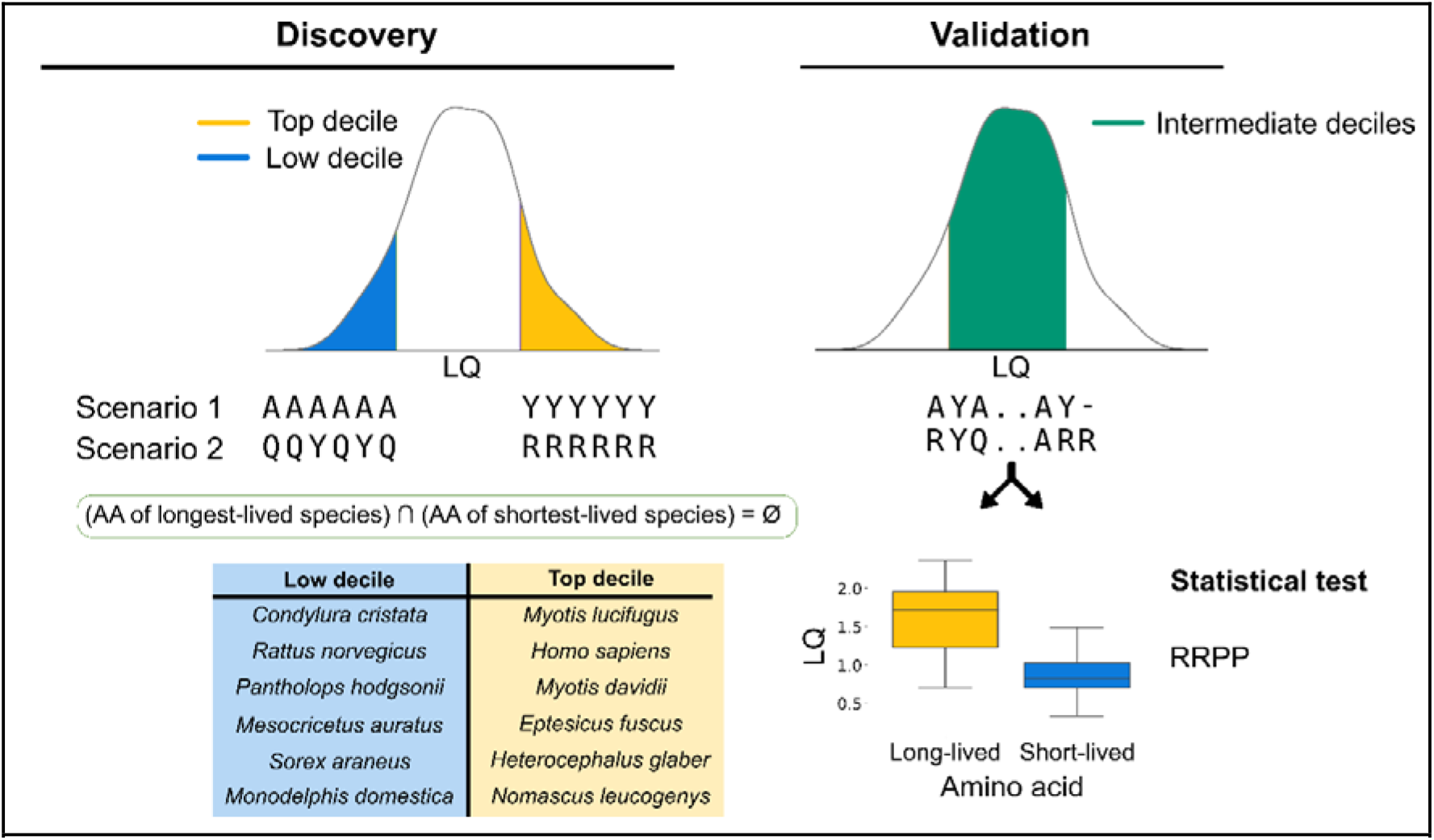
Workflow used in this study for the detection of Convergent Amino-Acid Substitutions (CAAS). In the Discovery phase, we identified AA substitutions that were exclusive from species in the top (yellow) and low (blue) deciles. In the Validation phase, we classified the species from intermediate deciles (red) in two groups, the species having the “long-lived” and the “short-lived” AA version. Finally, we ran a RRPP phylogenetic ANOVA to validate each discovered AA, keeping as significant only those for which we validated the direction of the effect (FDR<0.05).

To ascertain whether the number of CAAS identified as linked to extremely-lived species groups was different than random expectations, we performed two resampling tests. In both tests, two groups of 6 species were randomly taken from the phylogeny 1,000 times and the procedure to identify CAAS was repeated. The p-value was the empirical probability of getting a number of CAAS equal or larger than the original observation. The two resampling procedures differed in their consideration of the phylogeny. The first resampling was independent of the phylogeny; the species were selected completely at random *(random* resampling). The second resampling method was designed to maintain the same proportion of species in each order as those in the observed data *(guided* resampling). For mammalian orders where there were no other species to resample, we were conservative and always included the same species.

The second phase, a *Validation phase*, was applied to each AA pin-pointed in the *Discovery phase*. It consisted in validating whether the species in the intermediate deciles (middle 80% of the LQ distribution, a total of 45 species) that had the same AA as the long-lived species also had a higher LQ than those species having the same AA change/s as the short-lived species. When short-lived species displayed more than one AA, all of them were included in the Validation phase. However, AA present among the species of the intermediate deciles but that were not observed in the long-lived or the short-lived group, were discarded. For validation, we used a phylogenetic ANOVA test as implemented in the *RRPP* package in R, using 10,000 iterations for significance testing (Collyer and Adams 2018, Figure 1). Finally, to further validate the longevity signal recovered in the gene set we also performed an external validation with another mammal set of species (Supplementary Note).

### Annotation of CAAS

We analyzed the functional effects of the nucleotide changes leading to CAAS, their population frequency in humans and their association to complex diseases. The most likely nucleotide substitutions corresponding to AA substitutions associated with high LQ in mammals were ascertained using the *panno* option from TransVar (Zhou et al. 2015) and visualized in the protein context (Supplementary Note). The frequency of each genetic variant in current human populations was obtained from the GnomAD v3 variant database (Karczewski et al. 2020). In those positions showing variable AA in the short-lived species (Scenario 2), we selected the more conservative option to avoid duplicated sites. That is, we assessed all possible combinations and kept the alternative with the highest allele frequency in humans. Also, those cases in which the most plausible variation leading to the AA mutation implied the change of more than one nucleotide of a codon were excluded from the variation analysis. The same procedure was repeated for 100 random sets of AA substitutions to test whether our observations on genetic variation in the discovered positions fitted the random expectations (Supplementary Note).

The functional prediction of the genetic variants, as well as the SIFT and PolyPhen2 scores were obtained from the Variant Effect Predictor (VEP, McLaren et al. 2016). SIFT and PolyPhen2 predict the functional impact of an AA substitution, the first by leveraging the sequence homology and physical properties of the AA (Ng and Henikoff 2003), and the later by using physical and comparative models based on evolutionary conservation and structure (Adzhubei et al. 2010). CADD scores were obtained from the CADD project website (https://cadd.gs.washington.edu/, Rentzsch et al. 2019), and used to assess the deleteriousness of genetic variants, by classifying those with a Phred score higher than 30 as likely deleterious variants.

For the identified positions in Scenario 1, ancestral states were reconstructed to assess the likelihood of the last common ancestor of mammals harboring the putatively long- or short-lived AA. Simulation of the ancestral AA was performed using an empirical Bayes method as implemented in the R package *phytools* (Revell 2012). To avoid cases in which the ancestral AA was uncertain, we only kept those in which the AA in the root of the tree had a probability higher than 0.8. We then quantified the cases in which the ancestral reconstructed AA was the one present in the short-lived or long-lived mammals. Additionally, for Scenario 1 substitutions, we simulated 100 stochastic character maps using a fixed transition matrix that assumes the same rate of change for any AA transition to estimate the number of AA changes of each type, in order to quantify the number of changes across the phylogeny from any AA to the long-lived or to the short-lived AA (Huelsenbeck, Nielsen, and Bollback 2003).

### Protein models

We aimed to compare protein energy variations between short- and long-lived mammals. Since *Rattus norvegicus* proteins have been the object of numerous studies, we selected that species as the representative of short-lived group of mammals in our analysis. Humans were selected as long-lived representatives. Not all proteins had known structures for both organisms. Out of 104 protein sequences in which we have discovered CAAS and that have known structures, we only found 40 protein sequence-pairs with high similarity and validated CAAS, from both human and rat. Then, we used MODELLER (Webb and Sali 2016) to model both structures of the pair, using the structures of the known templates and the sequence alignments obtained with “matcher” (from EMBOSS package) (Rice, Longden, and Bleasby 2000). The modelled structures were optimized with the repair-pdb protocol from FoldX (Buß, Rudat, and Ochsenreither 2018). We used the optimized structures to calculate the differences of Fold X energies (ΔE) between the human and rat protein sequences.

We selected all the sequences of *Rattus norvegicus* available in Uniprot with known 3D structure (n=667) as a background set to compare the ΔE distributions of proteins coded by genes harboring CAAs with the distributions of ΔEs of pairs of sequences from genes without CAAS. After removing the ones in the set with CAAS described in the previous paragraph, we obtained 337 structural models of highly similar human sequences. Note that we were not able to model the structure of all proteins due to threading mismatches. Outliers were removed from both distributions by generating an Interquartile Range (IQR) with a weigh of 4: IQR = 4*(upper quartile - lower quartile). The distribution was normalized by the maximum value to have comparable ranges between 0 and 1. We compared the distribution of the pairs with validated CAAS with the background distribution of non-validated CAAS using a permutation two-sample test (https://statlab.github.io/permute/user/two-sample.html).

We further tested the robustness of the results by increasing the size of the background. We increased the background with protein-pairs of highly similar human and rat sequences without validated CAAS, selected randomly. After parsing around 4,000 sequences, we obtained 500 pairs of rat-human sequences whose structure could be modelled for both species. We used the same protocol for modelling and optimization of the structures, and the distribution of ΔEs was similarly normalized and analyzed, using a permutation two-sample test for the comparison.

### Gene-Phenotype Coevolution across mammals

The nucleotide alignments that underwent previous quality control (< 50% of gaps) were used for studying the coevolution of genes and phenotype. We estimated root-to-tip rates of protein evolution (the dN/dS ratio or □) using the free-ratio model from PAML 4.9a (Yang 1997). The root-to-tip ω is a property of the species tip rather than of the terminal branch, thus being more inclusive of the evolutionary history of a locus and, therefore, it is more suitable for regressions against phenotypic data from extant species (Montgomery and Mundy 2012). Briefly, for each gene and species we computed the root-to-tip dNs and dSs and the ratio between these values to obtain the root-to-tip □, as previously described in Muntané et al. 2018. To avoid numerical problems with the log transformation and unrealistic substitution rates, the species for which the root-to-tip ω was 0 were discarded, with 877 genes having at least one species removed. Genes for which we could not estimate a root-to-tip value for, at least, half of the species were also removed from further analyses, resulting in a final set of 17,969 genes.

For each gene, we studied the association between its rate of protein evolution and longevity regressing root-to-tip ω and LQ by means of phylogenetic generalized least squares (PGLS) as implemented in the *caper* library in R (Orme 2018). PGLS allows to incorporate the phylogenetic relationship among species in the error term of a generalized least squares model, thus controlling for the phylogenetic inertia (close species may have more similar phenotypes than distant species). Pagel’s lambda (λ) was estimated through maximum likelihood in each case. Pagel’s λ values equal or close to 1 indicate that a character is evolving stochastically (Brownian motion) along the tree, whereas λ□<□1 indicates that a character evolution is independent of the phylogeny. In 295 genes, estimations of λ resulted in a value of 0 and the log-likelihood plots showed a flat likelihood surface (for an example, see Supplementary Figure 2), which is most likely due to reduced sample size (DeCasien, Williams, and Higham 2017). Consequently, for these cases we set the value of λ to the genome-wide median λ value. To control for the effect of effective population size covarying with LQ, median genome-wide root-to-tip □s was included in the PGLS models as covariate (Boddy et al. 2017). Also, species with studentized residuals > ± 3 were considered outliers and, thus, removed from the regression and PGLS was fitted again. Moreover, to control that no single species was biasing the PGLS models, we performed an additional step and repeated regressions: by removing one species at a time and keeping the maximum p-value (p-value conservative). In all regressions, both the LQ and the root-to-tip □s were log10 transformed. Finally, we applied a Benjamini-Hochberg False Discovery Rate (FDR) with an FDR of 5% for multiple test corrections. To avoid associations due to one single species, when we refer to genes that are nominally significant in the PGLS analysis, we always refer to those that were nominally significant for the p-value conservative regressions (P_cons_ < 0.05).

### Functional enrichments

To study whether there was an over- or under-representation of genes previously related to aging in our gene sets we used hypergeometric tests. We included lists of genes that had been previously associated with aging (Supplementary Note). Biological mechanisms underlying CAAS were evaluated with WebGestalt, that allows checking for pathway over-representation of specific GO terms, pathways, and disease-associated genes from GLAD4U (Liao et al. 2019). For each evaluated gene set, FDR was controlled using the Benjamini–Hochberg procedure and the set of evaluated genes (n=13,035 genes) was used as the background. PGLS results were filtered keeping only those genes that were nominally significant (P_cons_ – 0.05) and then ranked by the t-statistic obtained in the PGLS analysis, subsequently Gene Set Enrichment Analysis (GSEA), from WebGestalt, was carried out to study enriched categories in genes with both a positive and a negative association between root-to-tip □ and LQ.

### Human lifespan GWAS heritability enrichment

To test whether the genes obtained from our comparative analyses could explain genetic variation in human lifespan, LD-score regression (LDSC) was used (Finucane et al. 2015). Specifically, we tested whether both, the genes with CAAS (discovered and validated) and those nominally significant after the PGLS regression, explained a larger fraction of SNP heritability in a human lifespan GWAS than would be expected by chance. Briefly, SNPs on the GWAS of parental lifespan (Timmers et al. 2019) were assigned to genes by annotating and keeping only SNPs in genic regions plus a window of 5kb around each gene. Subsequently, the custom annotation file for LDSC was prepared for five categories: (i) all genic SNPs, (ii) SNPs that map into the genes that were evaluated, (iii) SNPs located in the genes containing discovered CAAS, and (iv) SNPs in the genes harboring phylogenetically validated CAAS and (v) SNPs in the genes that were significant (P_cons_<0.05) in the PGLS genome-phenome analysis. Enrichment in human lifespan heritability was evaluated in the detected genes (cat. iii, iv and v) compared heritability explained by the genes evaluated (cat. ii). To help in visualizing the results, stratified QQ-plots were also performed with the SNPs in the five categories.

Pathway Scoring Algorithm (PASCAL) is a software that allows testing if a given gene set (or pathway) is enriched in GWAS signal (Lamparter et al. 2016). With this aim, we computed gene scores by aggregating, using the sum of chi-squared option (SOCS), SNP p-values from the GWAS on parental lifespan (Timmers et al. 2019), while correcting for linkage disequilibrium data. These computed gene-based scores were then aggregated across sets of related genes with the pathway analysis tools in PASCAL to obtain a pathway score. Pathway enrichment was evaluated using the chi-squared method. With the aim of testing whether our identified genes were enriched in human lifespan GWAS signal, we built custom pathways using the genes resulting from our analyses (the five categories aforementioned) and computed pathway scores for them. For all of them, we kept only genic regions including a window of 5kb around each gene.

## Results

### Convergent Amino Acid Substitutions: Discovery and Validation

To detect convergent AA substitutions (CAAS) shared between long-lived mammals, we split species into two groups selecting those in the extreme deciles of the LQ distribution (rounding up to 6 species in each group, Supplementary Figure 1). Throughout the manuscript we used the term “Convergent” rather than “Parallel” AA Substitutions to acknowledge that we cannot guarantee that each substitution had appeared independently in each species. In a first phase, the Discovery phase, we counted all AA changes in which the same AA was present in the reference genomes of the long-lived species, while the short-lived species presented either (i) a different fixed AA (Scenario 1) or (ii) variable AA different from the one in the long-lived group (Scenario 2). The species included in the Discovery phase were three Chiroptera *(Myotis lucifugus, Myotis davidii*, and *Eptesicus fuscus)*, one Rodentia *(Heterocephalus glaber)*, and two Primates *(Homo sapiens* and *Nomascus leucogenys)* in the long-lived group, and two Soricomorpha *(Condylura cristata* and *Sorex araneus)*, two Rodentia *(Rattus norvegicus* and *Mesocricetus auratus)*, one Didelphimorphia *(Monodelphis domestica)*, and one Artiodactyla *(Pantholops hodgsonii)* in the short-lived group (See Figure 1 and Supplementary Table 1). Since gaps were not accepted in any of the sequences, the number of genes that were screened for the CAAS discovery was reduced to 13,035. We scanned all the aligned positions finding a total of 2,737 CAAS in 2,004 genes: 284 belonged to Scenario 1, and 2,453 to Scenario 2. We also identified 533 CAAS belonging to Scenario 3 (Table 1). It is worth noting that CAAS discovered in Scenario 2 were 4.6 times more frequent than CAAS in Scenario 3. If AA changes occur at random, the difference in the number of discoveries in Scenarios 2 and 3 is very unlikely (chi-squared P=1.88e-270), which constitutes strong evidence for an evolutionary trend to increased lifespan in the mammalian lineage.

**Table 1.** Lists of discovered and validated CAAS and genes. Numbers in parentheses represent the percentage of phylogenetically validated positions.

|  | Discovered |  | Validated RRPP |  |
| --- | --- | --- | --- | --- |
|  | CAAS | Genes | CAAS | Genes |
| **Scenario 1** | 284 | 273 | 131 (46.1%) | 128 |
| **Scenario 2** | 2453 | 1822 | 1025 (41.8%) | 891 |
| **Scenario 3** | 533 | 495 | 185 (33.5%) | 182 |
| **Scenarios 1 + 2** | 2737 | 2004 | 1157 (42.3%) | 996 |

We performed two different resampling tests to evaluate if the number of detected CAAS was higher than expected by chance: either randomizing species independently of their phylogenetic relation or randomizing only within mammalian orders (*random* and *guided* resampling, see Methods). The probability of randomly obtaining a number of CAAS equal or higher than the observed one was 0.003 in both resampling tests (Supplementary Figure 3), showing that our set of genes contains a statistical excess of CAAS and that, it is enriched with AA substitutions and genes linked to mammalian longevity after correcting by body mass. In our dataset, the *Didelphimorphia* and *Soricomorpha* orders only contained two and one species, respectively, and so the same species (i.e., *Monodelphis domestica, Condylura cristata* and *Sorex araneus*) were always included in the *guided* resampling, which resulted in a conservative test.

In the Validation phase, we confirmed the discovered CAAS using the species in the intermediate deciles of the LQ distribution. First, we divided them into two groups, those with the same AA as the long-lived species and those with the same AA combination as the short-lived species (Figure 1). Second, we tested whether the group presenting the long-lived AA showed significantly higher LQ using the phylogenetic ANOVA implemented in RRPP. Out of the 2,737 CAAS from Scenarios 1 and 2, we validated a total 1,157 that belong to 996 genes (Table 1 and Supplementary Figure 4). This resulted in a 42.1% validation out of the discovered CAAS (for a discussion on discovery and validation at other thresholds see Supplementary Note). Also, out of the 533 CAAS in Scenario 3, we validated 185 (34.7%) that belong to 182 genes. The list of all discovered and validated genes can be found in Supplementary Table 2. We should point out that for some cases the validation test was underpowered, as for some comparisons there were too few species in one of the two groups, which makes it even more important to validate such a high percentage.

Ancestral state reconstructions of the CAAS from Scenario 1 showed that, out of the 88 instances that were predicted with > 80% probability (Supplementary Table 3), only in four cases the long-lived AA was the ancestral state, while in the remaining 84 cases, the ancestral AA was the short-lived one (for an example see Supplementary Figure 5). To estimate the number of parallel AA changes in each gene from Scenario 1, we simulated 100 stochastic character maps for each AA substitution in the tree (a total of 28,400 simulations). We observed that in 79 out of the 284 AA substitutions from Scenario 1 there was an average of less than one change from any AA to the short-lived version, which implies that the short-lived AA was at the root of the tree. In 189 out of the 284 CAAS we observed less than 6 changes from any AA to the short-lived AA, while in 213 of the 284 CAAS, we observed less than 6 changes from any AA to the long-lived version, showing that the vast majority of CAAS appeared in parallel (Supplementary Figure 6).

We found that amongst the 2,004 discovered genes there was an enrichment of genes upregulated with age (FDR=9.43e-04, Enrichment Ratio (ER) = 1.43) and a depletion of age-downregulated genes (depletion FDR=9.99e-09, ER = 0.51), loss of proteostasis (FDR=1.05e-07, ER=0.26), essential genes (FDR=2.76e-06E-06, ER=0.72), and genes with pLI>0.9 (FDR=5.77e-19, ER=0.63). In contrast, there was no enrichment of genes previously associated with longevity from the GenAge database (Supplementary Table 4). The significant enrichments were conserved in the subset of 996 genes phylogenetically validated (Supplementary Note). Additionally, we studied functional enrichments in both the discovered and validated gene sets with WebGestalt (for a complete list of processes enrichments see Supplementary Table 5). The discovered genes were enriched in GO categories such as acute inflammatory response (FDR=1.99e-03), leukocyte migration (FDR=2.75e-02), and cytokine binding (FDR=2.96e-05), in pathways such as Staphylococcus aureus infection (FDR=6.34e-04), and complement and coagulation cascades (FDR=2.72e-03), and human diseases such as gram-negative bacterial infections (FDR=2.09e-07), autoimmune diseases (FDR=7.14e-06), systemic inflammatory response syndrome (FDR=1.17e-04), and Werner Syndrome (FDR=1.85e-03), among many others. Also, we found enrichment in hallmark gene sets (Liberzon et al. 2015) such as IL6 STAT3 signaling during acute phase response (FDR=3.70e-05) and blood coagulation cascade (FDR=9.50e-05).

Among the 2,004 genes harboring CAAS we found eight genes that have been previously linked with longevity. One example is the *WRN* gene, which plays a critical role in repairing damaged DNA, showing two mutations that differ between long-lived and short-lived mammals. One was from Scenario 1, with two AA clearly differentiating long- and short-live mammals (F1018L) and another from Scenario 2 (N1055S/R/K/I/T). Both mutations were in the RQC domain of the protein, which makes these two mutations good candidates for follow-up studies and experimental validation. Another example is *CASP10*, a gene involved in the activation cascade of caspases responsible for apoptosis execution, showed 6 CAAS, all of them located in the caspase domain and validated with the phylogenetic test (Supplementary Figure 7). A final example, *ZC3HC1*, which has been recently identified in the GWAS of parental lifespan (Timmers et al. 2019), contained a validated Scenario 2 substitution (T366S/A).

### Human variation in CAAS

Translating CAAS nucleotide changes from short-lived mammals to humans using TransVar, we found 2,704 out of the 2,737 CAAS mapped to a single nucleotide substitution. We excluded the remaining 33 CAAS because they needed more than one nucleotide substitution. We identified human genetic variation in only 516 out of the 2,704 CAAS (19%), but only in 6 cases (0.22%) the minor allele frequency (MAF) was higher than 1% (Supplementary Table 6). This suggests that the vast majority of the identified AA substitutions are fixed or almost fixed (99.78%) in humans and, thus, that they may correspond to genomic factors contributing to lifespan or related traits that are invisible to analyses that exploit variation in current human populations (e.g., GWAS). This observation is much lower than that expected by randomly selecting 100 subsets of 2,737 AA substitutions among the substituted positions between human and rat, and between human and green monkey (empirical P<0.01 in both). In the randomization we observed a mean percentage of 22.4%and 32.85% human genetic variation in the 100 subsets, respectively, and in all simulations the percentage was higher than the observed 19%. Moreover, the number of nucleotide substitutions with a MAF higher than 1% exhibited a mean of 0.60% and 2.36% and only in one out of the 100 randomizations between human and rat, the mean was lower than the 0.22% of the observed (Supplementary Figure 8).

Out of the 2,704 CAAS that mapped to a single nucleotide substitution, SIFT and PolyPhen information was obtained for 2,175. A total of 2,134 and 2,112 of the substitutions were considered benign or tolerated in humans according to PolyPhen and SIFT scores, respectively, with 2,079 being considered benign by both metrics. CADD scores evaluated the 516 variants showing human variation as likely benign (summarized in Supplementary Table 7). This represented, for all the scores, an enrichment of tolerated substitutions compared to 100 random samplings (empirical P<0.01, Supplementary Note).

### Protein models

We compared the FoldX changes of total energy between modelled structures of human and rat sequences in 40 protein pairs with validated CAAS. The difference of energy showed that the genes harboring CAAS code for proteins that are more stable in long-lived mammals (represented by human) than in short-lived organisms (represented by rat). To test whether this trend is general or is a property of longevity-related proteins, we analyzed the energies of all the sequences of *Rattus norvegicus* with known structure and without validated CAAS that had similar human sequences whose structure was either known or could be modelled. The permutation test proved the over stability of human sequences with a P-value of 6.3e-04 (Supplementary Figure 9A). This trend was further validated in a larger background set of about 500 structural models of human and rat sequences, and the significance was preserved with P-value of 4.5e-04 (Supplementary Figure 9B). In short: the accumulation of longevity-related differences in AA residues between short- and long-lived mammals has resulted in increased stability in these proteins in long-lived organisms. The specific role of these AA residues is unclear, as the variability of local energies of FoldX are not remarkable for any specific partial energy (with the only exception of Van Der Waals clashes).

### Gene-Phenotype coevolution

To identify genes with rates of protein evolution associated with changes in LQ across the mammalian phylogeny, we computed the root-to-tip □ for each gene and species and evaluated its association with LQ using PGLS. Among the 18,266 gene alignments, 297 were removed because for more than half of the species we were not able to compute a root-to-tip dN/dS, finally evaluating 17,969 protein coding sequences.

After FDR correction, in the PGLS analysis, four genes showed a significant association between gene root-to-tip □ and species LQ (Figure 2): *SPAG16* (P=3.58e-7, slope=3.14), *TOR2A* (P = 2.26e-7, slope=-2.44), *ADCY7* (P = 1.63e-06, slope=-2.79), and *CDK12* (P = 7.81e-06, slope=3.92). Among the 4 significant genes, two *(SPAG16* and *CDK12)* showed a positive association between rate of protein evolution and LQ, and the other two showed a negative association *(TOR2A* and *ADCY7)*. These associations between the root-to-tip □ and the LQ values in the four genes were strong, since even after applying the p-value conservative method (see Methods), they were still the top four genes in the analysis (Supplementary Table 8). Moreover, 705 genes showed a nominal significant association between rate of protein evolution and LQ (P_cons_<0.05).

**Figure 2.**
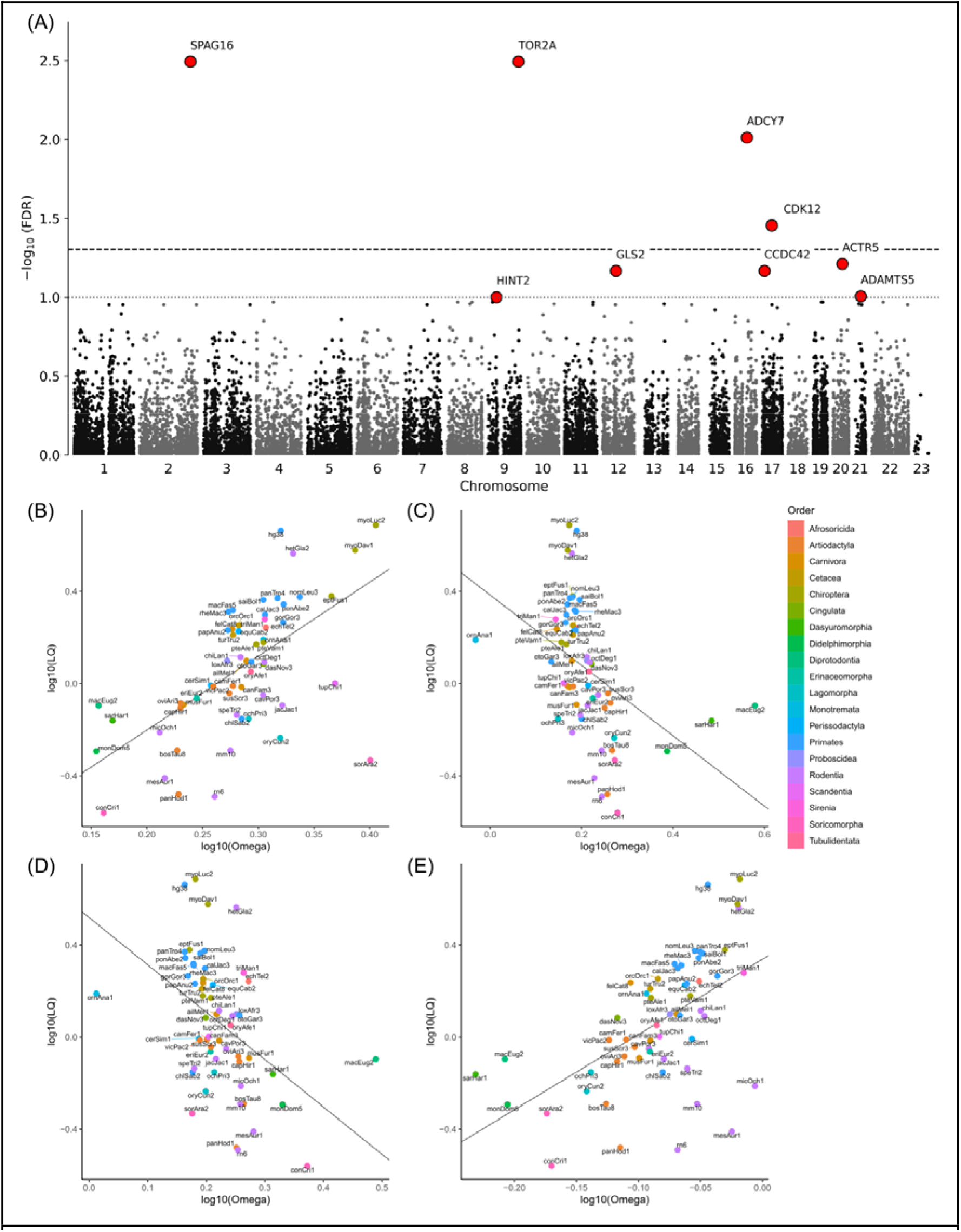
(A) Manhattan plot of gene-based association results of the phylogenetically-controlled regression (PGLS) for LQ. Each dot represents a gene and those depicted in red represent significance at FDR<0.1. The negative logarithm of the FDR P-value for each gene tested is reported on the y axis. P-value cutoffs corresponding to the Benjamini-Hochberg threshold FDR=0.05 and FDR=0.1, based on the 17,969 genes tested, are denoted by the dashed and dotted lines respectively. (B-E) Phylogenetically controlled regression (PGLS) between log10 root-to-tip ω for the significant genes (B) SPAG16, (C) TOR2A, (D) ADCY7, and (E) CDK12 are displayed against log10 LQ. Black lines represent the regression slope of the linear model. UCSC version names were used for species labeling. Correspondence to the species names can be found in Supplementary Table 1.

### Human lifespan GWAS signal enrichment

Finally, we evaluated whether the gene sets obtained in our analyses were enriched in current human lifespan array heritability as estimated from GWAS data. We used data from the largest, UK Biobank-based, study on human parental lifespan GWAS to date (Timmers et al. 2019). We partitioned heritability on the genic fraction of the SNPs using LDSC and observed a 3.4-fold enrichment of explained heritability in the set of genes with CAAS compared to the set of screened genes (P=3.46e-04). The enrichment was a 2.6-fold for the phylogenetically validated set (P=0.06). While genes with a nominal significant association (P_cons_<0.05) between rates of protein evolution and LQ showed a 4.2-fold enrichment in GWAS heritability (P=0.04, Supplementary Table 9). Figure 3 shows these enrichments using a stratified Q-Q plot, in which a leftward deflection from the null expectation of the subset of SNPs of interest implies an enrichment in GWAS signal. SNPs in the genes that were screened by the CAAS method did not significantly deviate from the expected p-values since it remained close to the line for all the SNPs from the GWAS. On the other hand, the discovered and validated genes, as well as genes that were nominally significant in the PGLS approach deviate from the null expectation and showed an enrichment on GWAS significant p-values. This enrichment on heritability from the parental lifespan GWAS was also confirmed using PASCAL (Lamparter et al. 2016). A chi-squared p-value of 3.27e-05 was obtained for the gene set comprising discovered genes with CAAS (Supplementary Table 10). The gene set created with the genes resulting from the phylogenetic validation also showed a significant enrichment (chi-squared P=0.039). Finally, the set of genes that were significant (P_cons_<0.05) after a PGLS between LQ and root-to-tip ω’s also showed significant enrichment (chi-squared P=8.66e-04).

**Figure 3.**
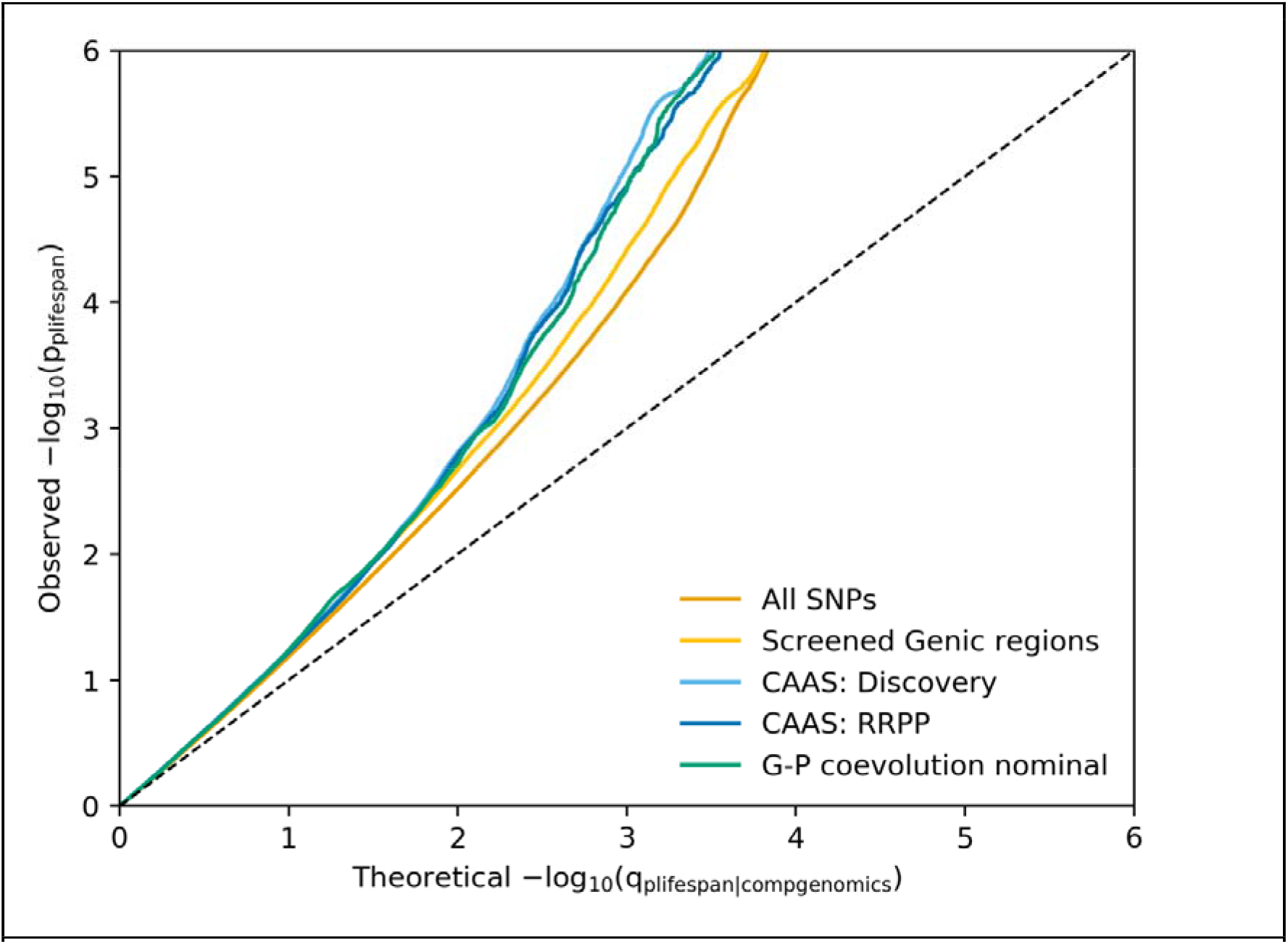
Stratified Q-Q plot for human longevity shows consistent enrichment across several assessed gene sets. Annotation categories were i) all SNPs in the GWAS (orange); ii) SNPs in genic regions of genes screened by the CAAS method (yellow); iii) SNPs in the discovered genes (light blue); iv) SNPs in genes validated with RRPP (dark blue); and v) SNPs in genes nominally significant (P_cons_<0.05) in the PGLS regression (green). All genic regions were defined by gene boundaries plus 5Kb. The genes we validated in the study were enriched in human longevity signal.

## Discussion

The largest-scale studies trying to unveil the genomic architecture of lifespan variation, including human GWAS, have focused on single species. As a consequence, these studies cannot detect variation that, while fundamental to define lifespan-related phenotypes, may have been fixed in the lineage of a species and may contribute crucially to differences in longevity across species. Comparative genome-phenome analysis, therefore, is essential to obtain a complete view of the genetic architecture of lifespan, to unveil important longevity-related genes and genomic features, and to understand the evolution of long-lived species. Here, we leverage longevity variation across mammalian species to explore cross-species variation and identify mutations and genes linked to the evolution of lifespan. The genes detected belong to pathways potentially involved in longevity, have an increased protein stability in long-lived species, and capture a significant part of the variance in the lifespan of current human populations explained by GWAS.

### Genetic architecture of longevity across mammals

Our comparative analysis discovered 3,270 Convergent Amino-Acid Substitutions (CAAS) in 2,314 genes using species in the extreme values of LQ distribution. Among them, 2,737 CAAS in 2,004 genes were from cases in which all reference genomes of long-lived species present the same reference AA, while short-lived species always present a different AA. This was a statistical excess of discoveries, emphasizing that our gene set is enriched with lifespan-related genetic variation. Out of the 2,737 discoveries, 1,157 CAAS (a 42.3%) in 996 genes were validated using the species in the intermediate deciles with a phylogenetic ANOVA test. The observed 42.3% is much higher than the 5% validation expected by chance, showing, again, that our approach can unveil true longevity signals. Furthermore, it should be noted that in the Validation phase, some of the CAAS could not be validated as there was insufficient statistical power due to the small number of species in one of the two groups.

Our results strongly support the use of a comparative genetics approach to inform and complement the interpretation of human lifespan GWAS. First, we observed the enrichment of GWAS signal in stratified QQ-plots of human lifespan. Second, focusing on genes that contain discovered and validated AAs, we evaluated whether the proportion of lifespan heritability that they explain was larger than expected by chance. Third, we analyzed a custom pathway created with the obtained gene sets for enrichment in GWAS by using PASCAL. All resulted in significant enrichments for the lists including genes from the convergent substitutions and the gene-phenotype coevolution analysis. Moreover, comparative genomics is the only way to pinpoint most of the genes we report here, since most of the detected AA changes were fixed or almost fixed in current human populations, with only 5 out of 2,230 substitutions (0.22 %) segregating at MAF > 1%. This is significantly less than what is expected by selecting random SNP across the genome and highlights the fact that variation associated with longevity in mammals is almost all fixed in humans. In sum, we demonstrated that an across-species comparative genomics approach can complement the analysis of the genetic architecture of complex traits like LQ.

A number of different biologically significant phenomena *(e.g.*, point mutations and post-translational modifications) can change the folding stability of a protein. Here, we found that the proteins harboring AA changes linked to increased lifespan show increased stability in humans compared to short-lived mammals, represented by rats. The exact cause of increased protein stability cannot be determined from the data collected in this work. Some trends suggest that over stability may be due to the contacts in the hydrophobic core, but results were not significant, with the exception of a reduction in van der Waals clashes. Still, an overall explanation for our findings may be that these proteins have accumulated AA changes resulting in increased resistant to the general proteome destabilization that comes with age. In fact, we also observed a significant depletion of genes linked to loss of proteostasis among the discovered gene set. These observations are consistent with evidence showing that long-lived animals have improved protein stability mechanisms (Kim et al. 2011; Pérez et al. 2009; Treaster et al. 2014). Taken together, this is compelling evidence that a common cross-mammalian mechanism to increase lifespan involves proteome stabilization.

In line with this observation, the identified longevity-related set was depleted in essential genes and genes intolerant of loss-of-function variation (pLI ≥ 0.9). Moreover, the deleteriousness scores of the mutations we discovered were enriched in tolerated substitutions compared to random mutations. This could be explained if lifespan evolves through genes and pathways that are not essential to the organisms, which would be fitting with the considerable evolutionary plasticity of longevity in mammals (Ratikainen and Kokko 2019).

It is of note that there were only 533 cases in which the long-lived species showed variable AA and the reference genomes in the short-lived species presented the same AA (Scenario 3). Assuming randomness, this represents a significant reduction from the 2,453 AAs in Scenario 2 (same AA in all long-lived species, different AAs in the short-lived ones). To the best of our knowledge, this is the first genomic evidence of a trend to increased lifespan in mammals, probably driven by positive selection throughout the mammal clade, which is consistent with the fact that stem mammals were small (O’Leary et al. 2013). This observation was validated by the fact that only a 4.5% of the long-lived variants from Scenario 1 were estimated to be present at the root of the mammalian phylogeny, while in the remaining 95.5% the mammalian ancestor was assigned the short-lived variants. In many cases, for instance, substitutions were present in a long-lived ancestor and are shared by sister species, even if there are no sister species in the top and bottom deciles used in the Discovery phase. However, for most AA changes in Scenario 1 we have been able to validate that long-lived species incorporated the AA mutation multiple times in parallel. These cases are examples of independent mutations that appeared in parallel with lifespan shifts across mammals, as described for other biological adaptations, such as echolocation (Liu et al. 2010) or adaptation to aquatic environments (Foote et al. 2015).

### Relevant genes and pathways

Discovered genes were enriched in processes involving immune and inflammatory response, cytokine binding and hemostasis, all of them pathways with well-known relationships with lifespan (Maynard et al. 2015). Coagulation, which has an important role in the maintenance of hemostasis, is known to increase with age and contributes to the higher incidence of cardiovascular diseases in the elderly (Franchini 2006; Khan et al. 2017). These pathways overlap with recent findings from Kowalczyk et al. 2020, where they used the number of AA substitutions on a phylogenetic branch to infer shifts associated with lifespan. They found that pathways such as inflammation, DNA repair, cell death, the IGF1 pathway, and immunity were under increased evolutionary constraint in large and long-lived mammals.

We did not find a specific enrichment in aging-associated lists among the uncovered genes. This might be explained by the fact that our analysis is intended to identify genes and pathways that are shared across mammals, while most of the aging-related genes discovered so far are mainly the result of single species approaches that capture genes driving current variation within species rather than crucial changes fixed along the phylogeny. However, many genes have been previously associated with longevity. A good example is two AA changes in *WRN* gene that were validated in our study at positions 1018 and 1055, both in the RQC domain, which is crucial for DNA binding and for many protein interactions (Tadokoro et al. 2012). *WRN* codifies for the Werner protein, which plays a critical role in maintaining the structure and integrity of the DNA. More than 60 mutations in the *WRN* gene are known to cause Werner’s syndrome, which is characterized by dramatically early appearance of features associated with aging. The two positions we identified (g.31141514T>C and g.31141706A>G) have not been reported before, because they show no variation in humans, suggesting a potential role of these specific positions in the evolution of lifespan in mammals.

Four genes showed a significant relationship between LQ and root-to-tip □ (PGLS at FDR<0.05): *TOR2A, ADCY7, CDK12* and *SPAG16*. Since they have co-evolved with longevity patterns across mammals, these are very good candidates for future aging studies. Some have been previously involved in regulating longevity-related pathways. For example, *TOR2A* is a gene involved in cardiovascular diseases (Sun et al. 2020) included in a CNV region in chromosome 9, correlating with longevity in a GWAS of Han Chinese individuals (Zhao et al. 2018). *ADCY7* is a key gene in the longevity regulating pathway and was recently associated with cancer mortality in dog breeds (Doherty et al. 2020). *CDK12* has been observed to be required for stress activated gene expression (Li et al. 2016). In contrast, SPAG16 has not been previously associated with longevity. Overall, using both approaches, we provide a list of genes that align with previous observations, but there are also new longevity-associated genes and mutations that will need to be validated experimentally.

### Limitations of the study

We should acknowledge some limitations in our study. First, we analyzed a reduced number of species and genes. Given the number of species and quality of gene alignments, we could get genetic and phenotypic data for 57 mammals. Out of 19,170 genes, we evaluated 13,035 good-quality genes. Very recently, the genomes of 250 mammalian species, including 132 assemblies, have been released (Zoonomia Consortium 2020), that resource can eventually be used for analyses similar to the one presented here. However, acquisition of high-quality samples was a major issue in that work (for instance, only 22 out of the 132 assemblies present contigs of N50 quality > 100Kbps) so the inclusion of these fragmented genomes would impact the comprehensiveness of our study because of a drastic reduction of the number of genes that we would be able to consistently analyze over all the phylogeny. Second, our power to validate CAAS depends on the numbers of non-extreme species showing either the long-lived or the short-lived amino acid residue. In some cases, the long/short-lived amino acid identified was not present in any other mammal, or only in a few, which made validation impossible. These cases are of course of interest, but beyond the scope of the present work. Third, because the decile strategy was decided *a priori*, it is likely that we have not explored the full set of AA changes that were involved in changes in mammalian lifespan. However, Supplementary Figure 2 suggests that using more extreme longevity values to classify CAAS may provide even more information of the genomic architecture of lifespan. Fourth, in this study we used the short-lived mammal with the largest available data on protein folding (rat and human), to demonstrate the level of stabilization of the proteins coded by aging-related genes detected in this study. However, the comparison was performed upon the reduced set of proteins with available folding data, which may not be representative of the whole proteome. Finally, in the CAAS method we identified point mutations that are ultimately assigned to genes. To study the heritability explained by these genes we assigned to each gene all the SNPs lying in its coding region (plus a 5kb window). This can introduce a bias because of complicated LD patterns and because many parts of the gene might not be relevant even if a GWAS association is lying in the gene. However, such bias would of course be conservative, even in cases where there is a GWAS signal lying in a gene.

### Conclusions

Our findings provide evidence on the genes and cellular mechanisms that may play a role in regulating mammalian lifespan, strongly suggesting that protein stability is linked to increased longevity and supporting an evolutionary trend towards longer lifespan in mammals. Moreover, our study is the first to showcase how comparative-genomic studies can illuminate the genetic architecture of human traits, including clinical and medical phenotypes, and supports the use of comparative genomics studies to understand complex human traits.

## Acknowledgements

We thank T. Marquès-Bonet for his insights discussing the analyses. This work was supported by: AEI-PGC2018-101927-BI00(FEDER/UE), the Spanish National Institute of Bioinformatics of the Instituto de Salud Carlos III (PT17/0009/0020), FEDER (Fondo Europeo de Desarrollo Regional)/FSE (Fondo Social Europeo), “Unidad de Excelencia María de Maeztu”, funded by the AEI (CEX2018-000792-M) and Secretaria d’Universitats i Recerca and CERCA Programme del Departament d’Economia i Coneixement de la Generalitat de Catalunya (GRC 2017 SGR 880).

