## Supplementary Note for "Comparative analysis of mammal genomes unveils key genomic variability for human lifespan"

### **Supplementary Methods**

**Discovery Phase: Scenarios 3 and 4**

For the purposes of this work, we focused on CAAS from Scenarios 1 and 2, corresponding to AA substitutions that were present in the reference genomes of the group of long-lived mammals. Our in-house script also includes two other scenarios: Scenario 3 and Scenario 4. Scenario 3 was defined as AA positions where the all the reference genomes of the short-lived group of mammals present the same AA, and where different, non-intersecting, AA variation was observed in the long-lived group (the reverse of Scenario 2); while Scenario 4 contained CAAS that were different within each group, long- and short-lived mammals, and non-overlapping between groups. Information from scenario 3 was kept for comparison with the number of discoveries with those from scenario 2; however, results from scenario 4 were discarded as they were too noisy and there was no statistical power for their individual validation.

**Visualization of the identified AA mutations**

Additionally, specific protein domains and features were retrieved from UniProt and the location of each identified AA change was displayed in protein diagrams with the Lollipop software (Jay and Brouwer 2016). The software produces a concise visualization of the number of CAAS per gene and allows ascertaining whether they were or not in an active domain.

**External validation**

We used an external set of 31 mammals to further validate our findings (Supplementary Table 11), all of them available at NCBI, except for the genomes of the Balaena mysticetus, which was downloaded from the bowhead whale genome resource (Keane et al. 2015). For each of the mammals, we downloaded their protein blast database, and for each of the CAAS detected in our analysis we performed a blastp of the human sequence using an e-value cut-off of 1e-5 and keeping the best hit. We then classified the species in two groups, those that have the long-lived AA, and those who have the short-lived AA and performed a phylogenetic ANOVA. Those CAAS in which we had less than 5 species in each group were excluded.

**Generating a random set of AA to check the expected percentage of variation in humans**

We checked if variation observed in the nucleotide changes leading to CAAS in current human populations was different than random expectations. To do so, we generated a set of AA positions by selecting, based on the same protein alignments, all the AA substitutions between human and rat, and human and green monkey. We used a set of positions within the universe of changing positions across mammals in the same genes in which we discovered CAAS in our analysis and excluded AA positions where no changes in mammalian evolution had occurred. Adding the latter would surely be biasing our AA set with positions that are non-variable, for instance due to strong purifying selection. Then, we randomly chose 100 subsets of same-length AA positions as the discovered CAAS in our analysis (n=2,737). For each AA substitution, we annotated the more likely nucleotide change leading to the AA substitution, using TransVar and the GnomAD variant database to identify human variation in those genomic positions. After 100 resamplings, we compared the percentages of variable positions and the positions showing genetic variation with MAF>1% in humans, against our observations.

For the same sampled substitutions, we tested whether the SIFT and PolyPhen information was different from our observation. We found that the lifespan-associated CAAS from our analysis were enriched in benign and tolerated amino acid substitutions. In our sample 97.1% and 98.1% of CAAS were predicted to be tolerated or benign according to SIFT and PolyPhen score respectively, while the mean percentage in the 100 subsets of AA substitutions between rat and human was of 94.5% tolerated (empirical P<0.01) and 97.8% benign (empirical P = 0.1881). The mean percentage in the 100 subsets of AA substitutions in green monkey was of 94.5% tolerated (empirical P<0.01) and 95.9% benign (empirical P<0.01).

**Gene lists used for testing pathway enrichments.**

To study genes that have been previously related to aging, several lists of curated human genes were obtained from different sources (Supplementary Table 12). We included three lists of genes: (1) a study of aging-related genes using Genotype-Tissue Expression Project (GTEx) data, identifying both, upregulated and downregulated age-associated genes (Jia et al. 2018); (2) genes linked to DNA damage repair (Pearl et al. 2015); and (3) the genes that have been linked to ageing in mammals available from GenAge (Tacutu et al. 2018).

We also evaluated whether our gene sets were enriched in genes that are intolerant to loss of function mutations comparing them against genes with a pLI > 0.9, coming from the gnomAD database (Karczewski et al. 2020). We also evaluated our gene sets against sets of essential genes from two studies: the first one identifyed genes that are cell-essential by inactivating genes at the DNA level in human cell lines ; while the second one identifyed the human orthologues of mouse essential genes (Georgi, Voight, and Bućan 2013). The first list of essential genes was used for discovery and the second one for validation of our results.

Additionally, we used gene ontology (GO) annotation, which describes how gene products behave in a cellular context, to select GO categories encompassing the mechanisms or functions corresponding to processes considered hallmarks of aging in López-Otín et al. (2013). Specifically, (i) Genomic Instability included genes in DNA repair (GO:0006281) and Nuclear lamina (GO:00005652) categories; (ii) Telomere Attrition included genes in Telomere maintenance (GO:0000723) and Telomere capping (GO:0016233); (iii) Epigenetic Alterations was constructed with genes from Histone modification (GO:0016570), DNA methylation (GO:0006306), and Chromatin remodeling (GO:0006338); (iv) Loss of Proteostasis included the Chaperone-mediated protein folding (GO:0061077), Autophagy (GO:0006914), and Ubiquitin-proteasome system (GO:0043161) categories; (v) Deregulated Nutrient Sensing contained Insulin receptor signaling Pathway (GO:0008286) and TOR signaling (GO:0031929); (vi) Mitochondrial Dysfunction was defined using genes from Response to ROS (GO:0000302) and Mitochondrial genome maintenance (GO:0000002) categories; (vii) Cellular senescence included genes from Cellular senescence (GO:0090398); (viii) Stem Cell Exhaustion was built with Stem cell proliferation (GO:0072089) category; and, finally, (ix) Altered Intracellular Communication was build using genes from Inflammatory response (GO:0006954) and Inflammasome complex (GO:0061702).

### **Supplementary Results**

**Discovery and Validation at other thresholds**

Given that the selection of the most extreme deciles was done *a priori*, we performed resampling using different numbers of extreme species in the *Discovery* phase and using the rest of species in the *Validation* phase (Supplementary Figure 4). Using three species at both extremes maximized the number of discovered CAAS (n=42,739) but reduced the percentage of validated CAAS to <35%. The number of validated genes peaked at 4 species in each group (44.9%) and maintained up to 6 species. When using 6 top and 6 low species, the number of discovered CAAS was smaller (n=2,737) but obtaining a good percentage of validated CAAS (42.5% of the discovered CAAS). Consequently, the setup including 6 species in each group (*i.e.*, top and low deciles) was considered suitable to reduce the number of false positives and maximize validation.

**External validation**

We could test 607 out of the 1,157 validated CAAS in the extra group of mammals, and 254 (41.8%) were nominally significant in the phylogenetic ANOVA, much more than the 5% that would be expected by chance, suggesting again that our list of genes was enriched in longevity signal (Supplementary Table 13). Moreover, among the non-internally-validated CAAS, 349 out of 818 (42.66%) were validated using this external set of mammal species.

**Functional Enrichments in the Validated set of CAAS**

We cross-checked the obtained gene set against other age-associated gene sets. Using Genotype-Tissue Expression Project (GTEx) data (Jia et al. 2018), we showed that amongst the RRPP-validated genes there was an overrepresentation of genes upregulated with age (on-tailed hypergeometric test, FDR = 1.25e-3, ER = 1.65), and a depletion of age-downregulated genes (depletion FDR = 1.93e-4, ER = 0.50). Additionally, there was non-significant overrepresentation of DNA damage response genes (P = 4.11e-2, ER = 1.51) (Pearl et al., 2015). In contrast, there is no overrepresentation of genes previously associated with longevity from the GenAge database (P = 0.44, ER = 1.05). Interestingly, amongst the RRPP-validated genes there was an underrepresentation of loss-of-function intolerant genes (pLI>0.9) (depletion FDR = 6.01e-5; ER = 0.73). Our gene set also showed a depletion of essential genes, (depletion FDR = 2.78e-2, ER = 0.78) (Supplementary Table 4).

Additionally, we studied functional enrichments in the validated gene set using WebGestalt. We found that the validated gene set was enriched in cytokine binding (FDR=1.22e-03), pattern recognition receptor activity (FDR=9.83e-03), which is crucial to initiate an innate immune response, and DNA replication (FDR=9.55-e06), among others (Supplementary Table 5).

Genes that were nominally significant with the PGLS analysis (n=705) were not enriched in any age-related curated gene list from literature but a significant depletion on pLI>0.9 genes (FDR=3.40e-03; ER=0.58, Supplementary Table 4). Those that showed negative correlations between the root-to-tip ω and LQ were enriched in pathways and GO processes involved in metabolism, such as copper homeostasis (WP3286, FDR 0.02), organic acid transport (GO:0015849, FDR=0.02); while those showing positive correlations presented a significant enrichment in tubulin binding genes (GO:0015631, FDR 0.03). The direction of the enrichments is given by the sign of the enrichment score (Supplementary Table 14).
