## Supplementary Figures for "Comparative analysis of mammal genomes unveils key genomic variability for human lifespan"

**Supplementary Figure 1.** Phylogenetic tree created using all the mammal species included in our study with Longevity Quotient (LQ) records from online databases (AnAge and Animal Diversity Web). Highlighted in blue, the 6 species with lower LQ values and included in the short-lived group for Discovery. Highlighted in yellow, the 6 species with higher LQ values and included in the log-lived group for Discovery. Lower values for LQ are shown in yellow; higher values are shown in blue (n = 57 species).

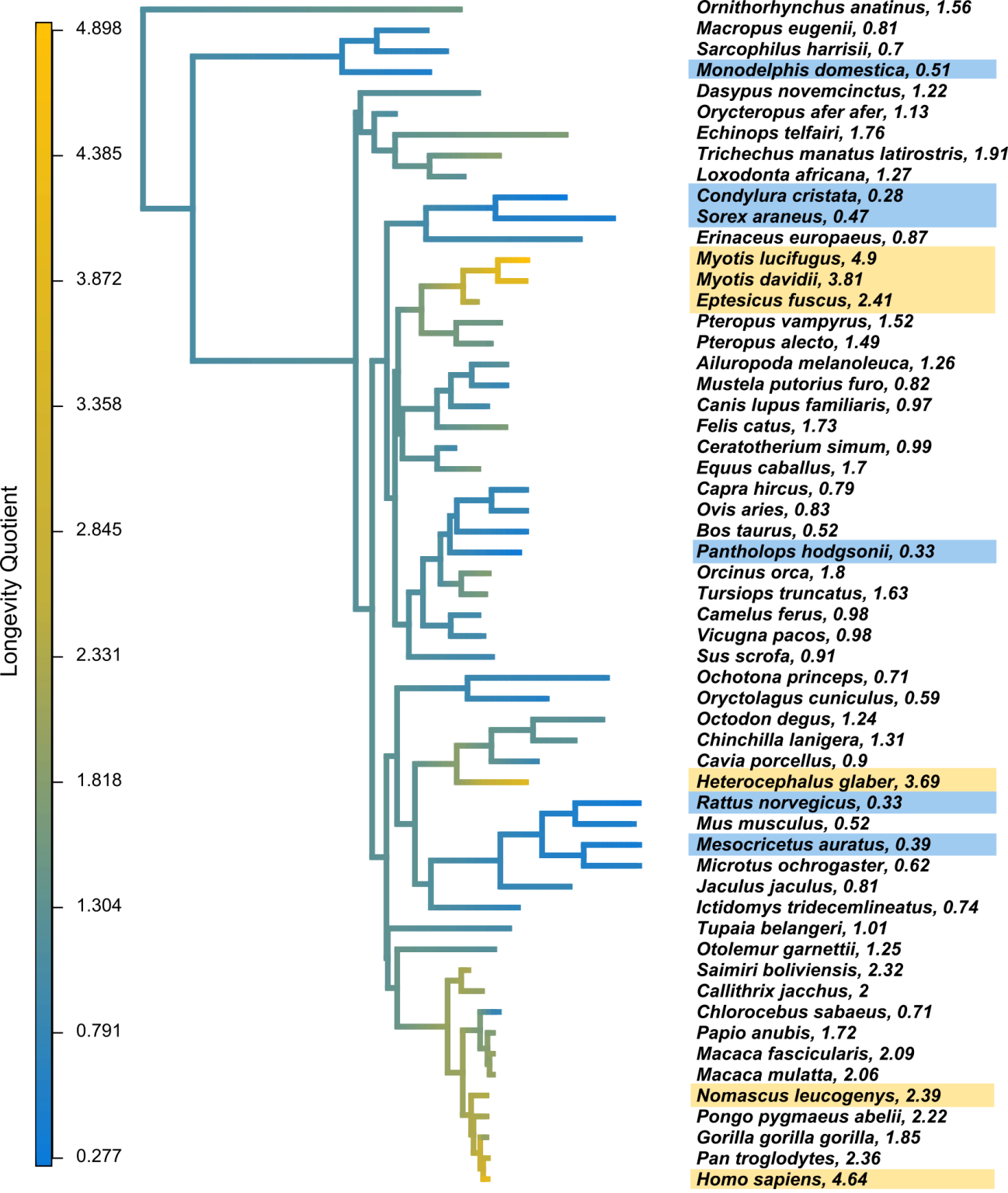

**Supplementary Figure 2. Lambda detection through maximum likelihood.** Representation of the lambda estimation for genes (A) *TOR2A*, and (B) *POLR3K,* where the maximum likelihood surface was too flat to properly estimate lambda.

| 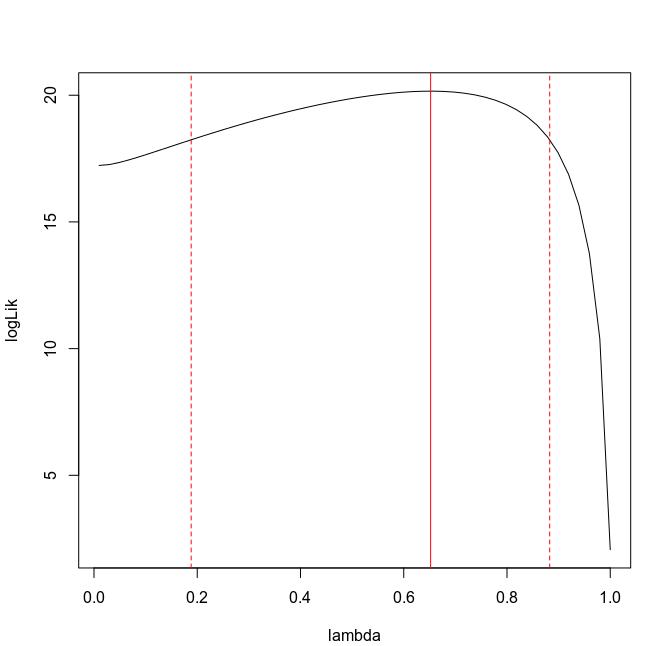 | 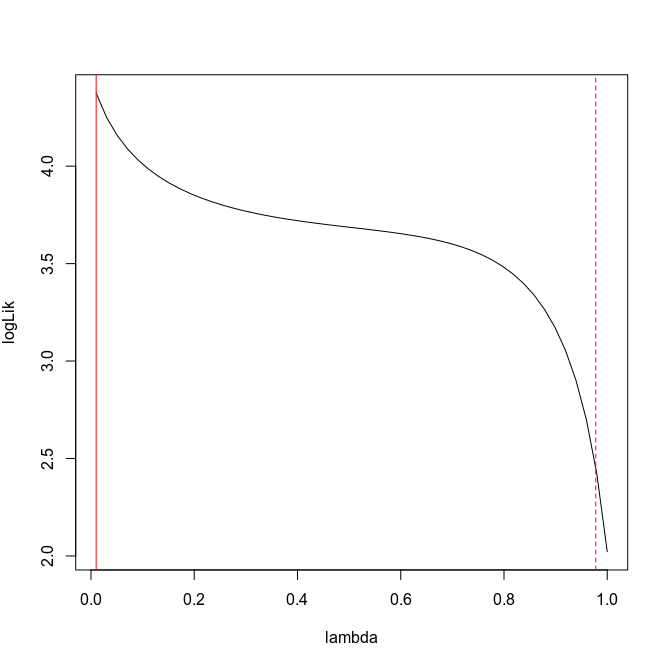 |
| --- | --- |

**Supplementary Figure 3.** Results of the resampling procedures (see Methods & Results). (A) random resampling; (B) guided resampling, keeping the proportion of orders in each group.

| A)  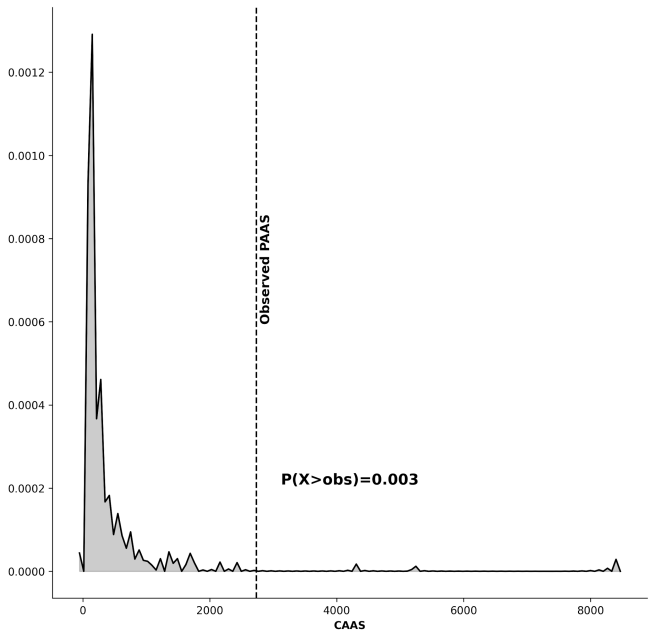 | B)  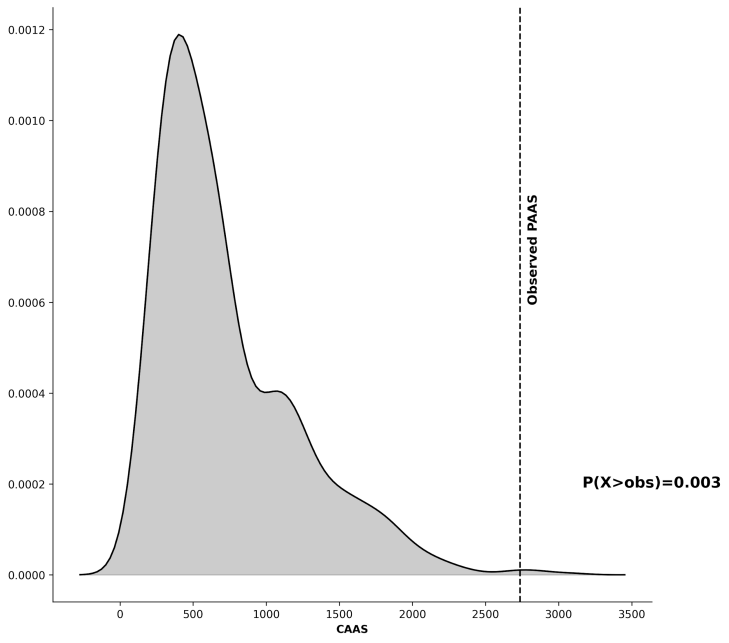 |
| --- | --- |

**Supplementary Figure 4. (**A) Bar plot of the number of discovered CAAS (left y-axis) when using different numbers of species (x-axis) for Discovery. The grey line represents the percentage of validated CAAS (right y-axis) using phylogenetic ANOVA (RRPP). (B) Table showing the 10 top long-lived species and the low short-lived species corresponding to the order of species that were used for Discovery. Species are sorted by descending LQ values in the long-lived column and by ascending LQ values in the short-lived column.

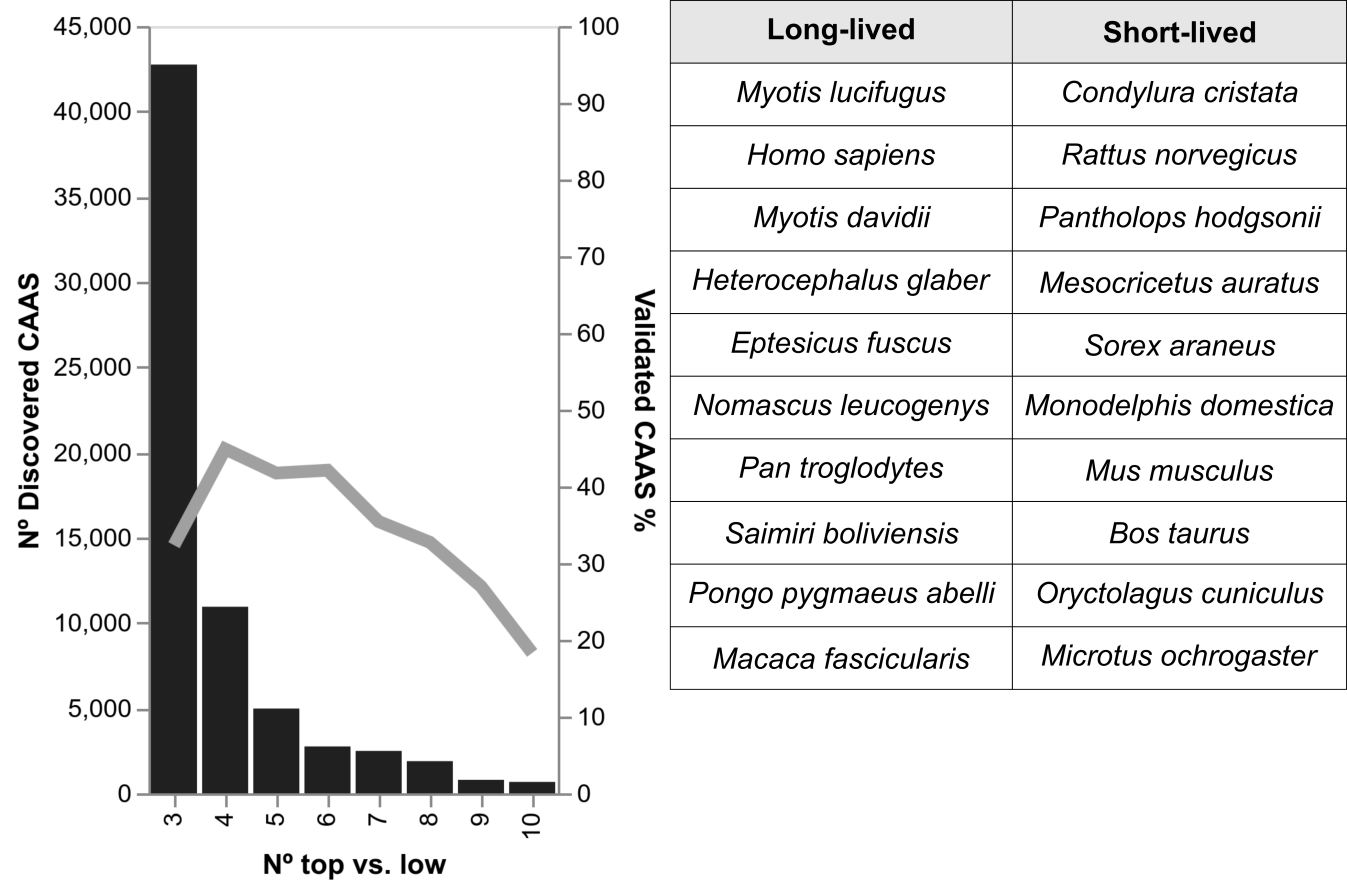

**Supplementary Figure 5. Ancestral reconstruction of PRUNE2 E1700K mutation.** AA position 1700 in *PRUNE2* gene is shown as an example. Note that the long-lived mammals carry a Glutamate (E in green), while the AA for the short-lived is a Lysine (K, in orange). The ancestral state was predicted to be a Lysine. The mutation to an E appeared in the branch leading to bats (*eptFus1, myoDav1 and myoLuc2*), in the naked-mole rat (*hetGla2*), and in the root of primates. However, the short-lived green monkey (*chlSab2*) carries the K mutation.

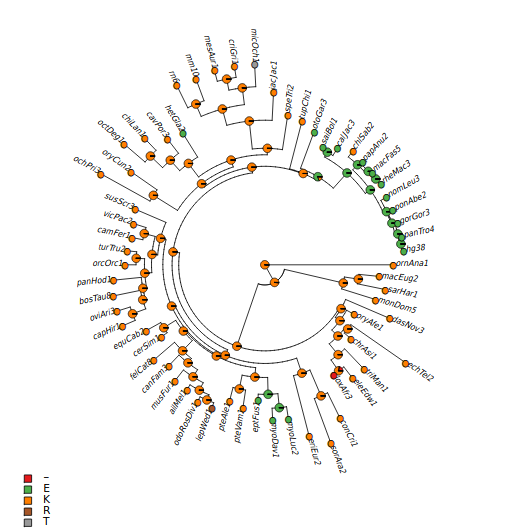

**Supplementary Figure 6. Distribution of estimated AA changes from any AA to the short-lived or long-lived one in the genes with CAAS from Scenario 1.** Note that we simulated 100 stochastic character maps for each CAAS, quantifying the mean number of changes from any AA to the short-lived AA, or the long-lived one.

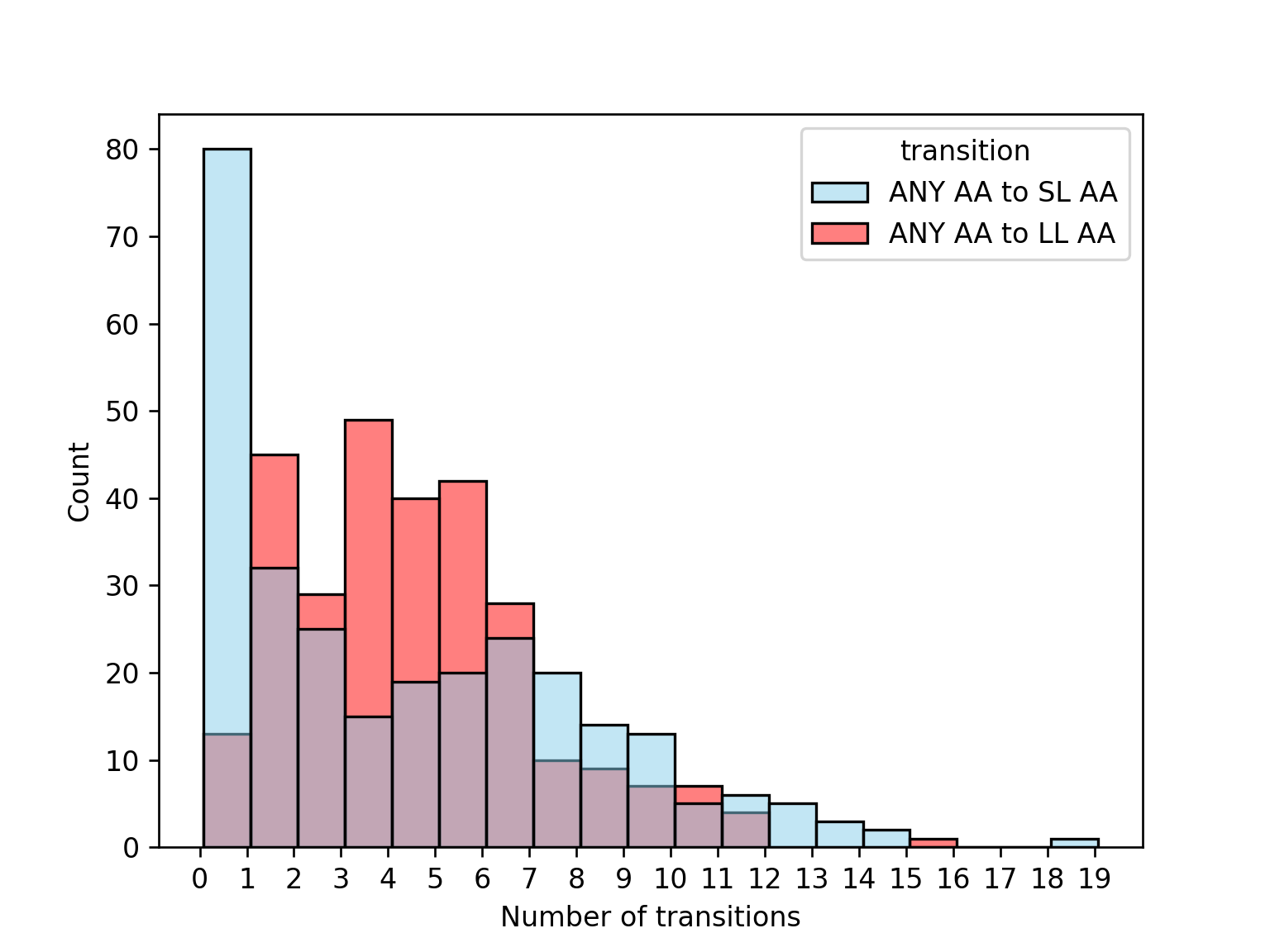

**Supplementary Figure 7.** Graphical representation of the CAAS found in the human proteins involved in lifespan A) *WRN*, B) *CASP10*, and C) *ZC3HC1*.

A)

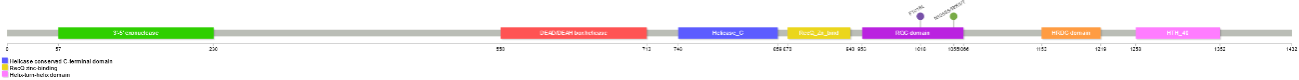

B)

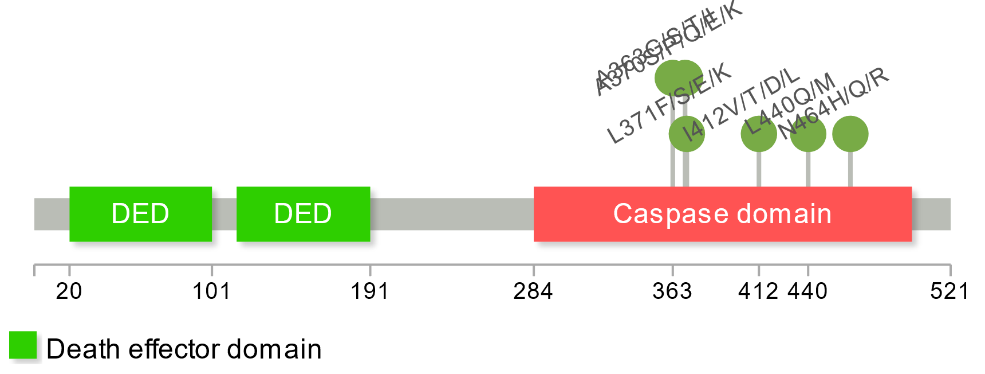

C)

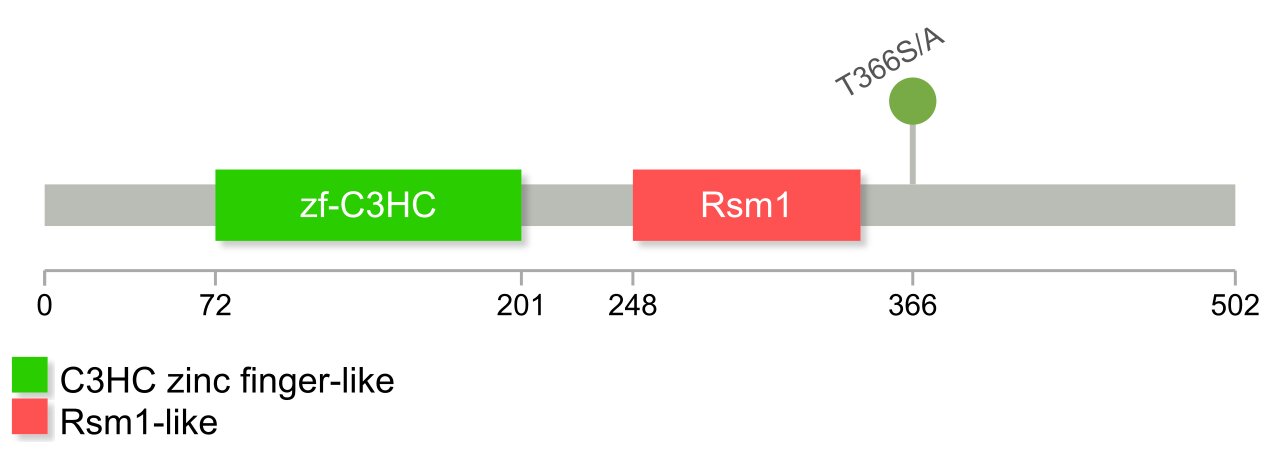

**Supplementary Figure 8.** Distribution of the percentage of human variation in 100 random subsets of AA substitutions, of the same size and in the same genes as our discovery set, between A) human and rat, and B) human and green monkey. Dashed line represents the real observation in our dataset.

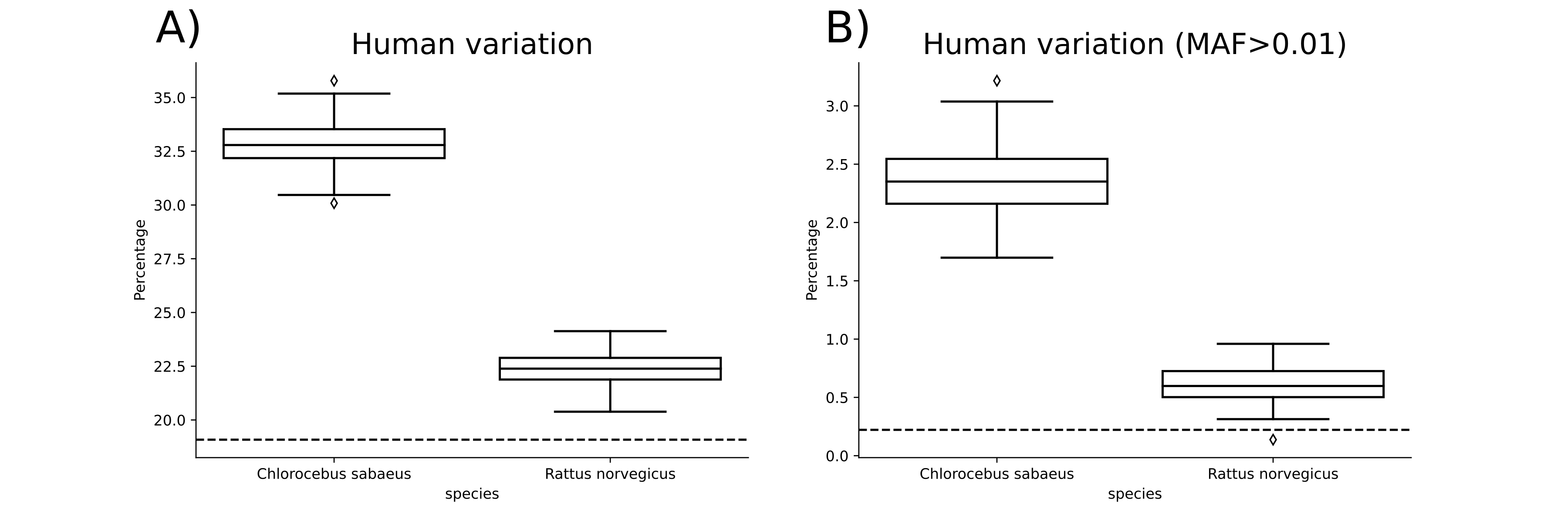

**Supplementary Figure 9.** Distributions of the difference of total energy (∆E), calculated with FoldX and structural models of rat and human sequences. A) Distribution for 40 validated CAAS (in red) compared against 337 non validated CAAS (in blue). B) Distribution for 40 validated CAAS compared against 500 non validated CAAS (in blue). Note: The number of proteins remaining after removing outliers is shown in the legend.

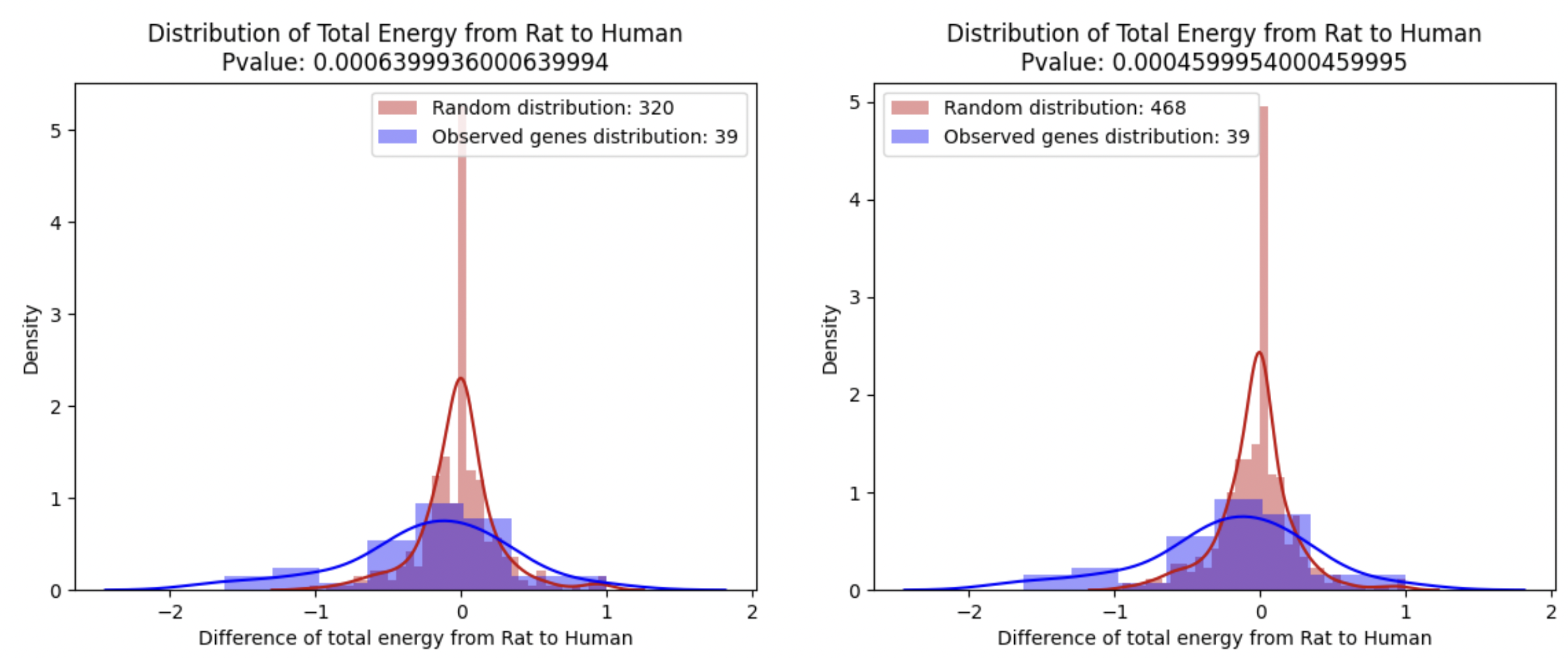
**A	B**
